# Cholesterol Dysregulation Drives Seed-Dependent Tau Aggregation in Patient Stem Cell-Derived Models of Tauopathy

**DOI:** 10.1101/2023.12.11.571147

**Authors:** Morrie Lam, Szu-Yu Kuo, Surya Reis, Jason E. Gestwicki, M. Catarina Silva, Stephen J. Haggarty

## Abstract

Tauopathies are a class of neurodegenerative diseases characterized by the progressive misfolding and accumulation of pathological tau protein in focal regions of the brain, leading to insidious neurodegeneration. Abnormalities in cholesterol metabolism and homeostasis have also been implicated in various neurodegenerative diseases. However, the connection between cholesterol dysregulation and tau pathology remains largely unknown. To model and measure the impact of cholesterol dysregulation on tau, we utilized a combination of *in vitro* and *ex vivo* tau aggregation assays using an engineered tau biosensor cell line and human induced pluripotent stem cell (iPSC)-derived neuronal cultures from an individual harboring an autosomal dominant P301L tau mutation and from a healthy control. We demonstrate that excess cholesterol esters lead to an increased rate of tau aggregation *in vitro* and an increase in seed-dependent insoluble tau aggregates detected in the biosensor line. We observed a strong correlation between cholesterol ester concentration and the presence of high-molecular-weight, oligomeric tau species. Importantly, in tauopathy patient iPSC-derived neurons harboring a P301L tau mutation with endogenous forms of misfolded tau, we show that acute dysregulation of cholesterol homeostasis through acute exposure to human plasma-purified cholesterol esters formed by the linkage of fatty acids to the hydroxyl group of cholesterol leads to the rapid accumulation of phosphorylated tau. Conversely, treatment with the same cholesterol esters pool did not lead to subsequent accumulation of phosphorylated tau in control iPSC-derived neurons. Finally, treatment with a heterobifunctional, small-molecule degrader designed to selectively engage and catalyze the ubiquitination and proteasomal degradation of aberrant tau species prevented cholesterol ester-induced aggregation of tau in the biosensor cell line in a Cereblon E3 ligase-dependent manner. Degrader treatment also restored the resiliency of tauopathy patient-derived neurons towards cholesterol ester-induced tau aggregation phenotypes. Taken together, our study supports a key role of cholesterol dysregulation in tau aggregation. Moreover, it provides further pre-clinical validation of the therapeutic strategy of targeted protein degradation with heterobifunctional tau degraders for blocking tau seeding.

## Introduction

The neuronal microtubule-associated protein tau encoded by the *MAPT* gene on Chr. 17 regulates microtubule stability, dynamics, and axonal transport (1, 2). Moreover, as an intrinsically disordered protein subjected to extensive post-translational modifications, tau can undergo rapid structural transitions to adopt different proteoforms playing various roles in signal transduction, regulation of synaptic functions, and DNA/RNA protection (3, 4). The misfolding and aggregation of tau into soluble oligomers and progressive formation of highly ordered β-sheet-rich fibrils, paired helical filaments (PHFs), and subsequently neurofibrillary tangles (NFTs) in neurons and glia of affected brain regions are the hallmarks of a class of neurodegenerative diseases known as tauopathies (5). These diseases can be sporadic or inherited through mutations in the *MAPT* gene (6).

To date, there are no effective disease-modifying or preventative therapies for tauopathies due in part to the incomplete understanding of the biological processes capable of driving tau misfolding and the complex molecular mechanisms leading to neuronal death. While the aggregation of tau into NFTs is a hallmark of tauopathies, the events leading to neuron loss and glial dysfunction may start much earlier. Growing evidence from cellular models (7, 8), murine tauopathy models (9), and postmortem human brain studies (10) suggest that low-molecular-weight, oligomeric forms of soluble tau may function as ‘seeds’ that act as the driver of higher-order tau assemblies and neuronal toxicity. The notion of pathogenic, oligomeric tau seeds is consistent with the observation that late-stage, insoluble tau tangles alone are insufficient to cause neuronal death in cellular and animal models of tauopathy (11, 12), along with the diversity of dysfunctional cellular phenotypes observed in tauopathy patient-derived iPSC models, including selective neuronal vulnerability to a range of cellular stressors, despite the absence of higher-order tau fibrils and tangles (13–16). Finally, there is some evidence that oligomeric tau ‘seeds’ can exhibit prion-like properties, as post-mortem patient-derived ‘seeds’ and recombinantly created ‘seeds’ can template the formation of tau aggregates both *in vitro* (10, 17, 18) and *in vivo* (8, 17).

Recently, growing evidence implicates cholesterol dysregulation as a driver of tau aggregation. Cholesterol homeostasis plays a critical role in preserving neuronal homeostasis as cholesterol is an essential building block of neuronal membranes and plays a critical role in the production and secretion of neurotransmitters, neuronal repair, membrane remodeling, and plasticity (19–21). In Alzheimer’s disease (AD), dysregulation of cholesterol homeostasis can alter amyloid precursor-protein processing and increase amyloid aggregation (22). Furthermore, altered cholesterol metabolism can impair synaptic vesicle exocytosis and reduce neuronal activity, leading to dendritic spine and synaptic degeneration (23, 24). Findings from van der Kant and colleagues have established a connection between cholesterol and tau aggregation, showing that cholesterol esters, the cellular storage form of cholesterol in which its hydroxyl group is linked through an ester linkage to the carboxylic acid moiety of fatty acids, can directly impair the proteasome, leading to tau accumulation in an *ex vivo* neuronal cell model of AD (25).

To model and study the impact of cholesterol dysregulation on tau aggregation in the context of a primary tauopathy, e.g., frontotemporal dementia (FTD), we utilized a multi-model approach to probe a causal link between tau misfolding/oligomerization and cholesterol homeostasis dysregulation. Our findings support a two-hit hypothesis indicating that cholesterol esters have the potential to expedite the progression of tau pathology once seed-capable, oligomeric tau species have been established. Furthermore, we employed a new chemical tool with specificity to misfolded tau, QC-01-175, a heterobifunctional small-molecule degrader capable of catalyzing the E3 ligase Cereblon (CRBN)-dependent ubiquitination and subsequent proteasomal degradation of oligomeric tau species (26, 27). Using QC-01-175, we demonstrated that dysregulation of cholesterol levels in neurons only impacts tau when in the presence of misfolded, disease-associated tau species and in a CRBN-dependent manner. Thus, QC-01-175 treatment effectively inhibited the tau pathology caused by exposure to excessive cholesterol esters. Taken together, our results support the notion that targeting misfolded tau species through proteasome-mediated degradation could be a viable strategy for preventing tau dysfunction caused by cholesterol dysregulation and, more broadly, for developing novel, early intervention therapies for age-related neurodegenerative diseases. Importantly, our results also support that targeting cholesterol homeostasis as a modifier pathway of tau aggregation might be a suitable therapeutic approach.

## Methods

### Recombinant Tau Purification

Recombinant Htau40ΔK280 was purified as previously described (28). In brief, htau40ΔK280, encoding the longest human Tau isoform with deleted K280 (kind gift from Mandelkow Lab, German Center for Neurodegenerative Diseases (DZNE), Bonn) was inserted into a pNG2 vector, a derivative of commercial vector pET3a (Novagen) containing an ampicillin resistance gene and expressed in BL21 (DE3) competent cells (Thermo Fischer, EC0114). Cells were grown in 1 L Luria broth containing 100 μg/mL ampicillin at 37°C shaking at 200 rpm until OD600 = 0.8. Tau expression was induced with 1 mM IPTG and expressed for four hours. Bacterial pellets were collected via centrifugation at 4°C and 5,000g and resuspended in ice-cold lysis buffer (50 mM MES pH 6.8, 1 mM EGTA, 0.2 mM MgCl2, 5 mM DTT, Roche complete protease inhibitor, 1 mg/mL lysozymes. 0.1% Triton-X). Cells were lysed by sonification, and the supernatant was collected via centrifugation at 4°C at 5,000 g. The supernatant was then supplemented with 500 mM NaCl and submerged in boiling water for 20 minutes.

Denatured proteins were pelleted by ultracentrifugation at 4°C and 200,000 g. The resulting supernatant containing soluble tau was collected and dialyzed against 20 mM MES, pH 6.8, 50 mM NaCl2, 1 mM EGTA, 1 mM MgCl2, 2 mM DTT, and 0.1 mM PMSF overnight. After dialysis, the supernatant was applied onto a HiTrap SP HP cation-exchange column (Cytiva, 17-1152-01), and tau was eluted using a 50 nM to 600 nM NaCl gradient. Tau positive fractions were collected and concentrated using an Amicon 10 kDa cutoff centrifugal tubes (Millipore, UFC5003) and gel filtrated in 30 mM Tris pH 7.5 and 2 mM DTT.

### Recombinant Tau Aggregation

To create tau aggregates, 50 μM recombinant tau was incubated with 8 μM heparin in 30 mM Tris pH 7.5 and 2 mM DTT at room temperature for 7 days. Every day, fresh 1 mM DTT was supplemented into the incubation. At the end of day 7, tau aggregates were pelleted by ultracentrifugation at 200,000g for one hour. Tau aggregates were resuspended in PBS, pH 7.4 with 2 mM DTT, and sonicated for 15 seconds at 30% amplitude and 250 Watts to create a homogenous population of tau fibrils. Aggregate concentration was measured by absorption at 280 nm using an extinction coefficient of 0.31 mL mg^-1^cm^-1^. Tau aggregates were diluted to desired concentrations in OptiMEM and incubated with lipofectamine 2000 (ThermoFischer, 11668019) according to the manufacturer’s instructions prior to treatments.

### Thioflavin-based Tau Aggregation Assay

The formation of tau aggregates was measured using the β-sheet binding dye thioflavin T (ThT) in a 96-well plate format based upon the principle that there is a redshift and increase in the fluorescence of ThT upon binding to amyloids (tau aggregates). Fluorescence measurements, with excitation at 450 nm and emission at 482 nm, were taken on an Envision 2102 Microplate Reader (PerkinElmer), with readouts taken every 20 minutes or hour with shaking at 37°C. In seeded reactions, 1 μg of recombinant tau aggregates was incubated with 20 μM recombinant monomeric tau in 30 mM Tris pH 7.5, 1 mM DTT, 20 μM ThT, and 5 μM heparin and respective LDL concentrations. In non-seeded reactions, 20 μM recombinant monomeric tau was incubated in 30 mM Tris pH 7.5, 1 mM DTT, 20 μM ThT, 5 μM heparin, and respective LDL concentrations. The background was corrected for where appropriate in the absence of recombinant tau aggregates and monomeric tau. The kinetic curves measured by ThT fluorescence were fitted to the following equation:

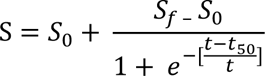

Where *S*_0_ is the absorbance signal at time zero, *S*_*f*_ is the final absorbance, t is time, and t50 is the time at which the change in signal is 50% the maximum.

### HEK293 GFP-tau Reporter Line Creation and Culture

Genes of interest were cloned into the pcDNA5/FRT expression vector and co-transfected with pOG44 into the HEK293 Flp-In T-Rex cell line (Thermo R78007) following the manufacturer’s instruction. Two days after the transfection, cells were subjected to selection with 150 µg/mL hygromycin and expanded for 2 weeks. Stable HEK293T cell lines with inducible expression of GFP-tau were grown in Dulbecco’s Modified Eagle’s medium (Gibco) supplemented with 10% fetal bovine serum and 0.1% penicillin/streptomycin at 37°C and 5% humidity. Growth media was supplemented with 150 µg/mL hygromycin every 3^rd^ passage.

### Generating Cereblon Knockouts in HEK293T GFP-tau Line

Synthetic guide RNA (sgRNAs) were designed to target the *CRBN* gene encoding Cereblon (Target DNA Sequence: CGCACCATACTGACTTCTTG, Target Locus: Chr.3: 3174104 - 3174126 on GRCh38) utilizing TrueCut^TM^ Cas9 V2 (ThermoFisher, A36498). HEK293T were seeded into 6-well plates at 250,000 cells/well the day prior to transfection. RNPs were delivered to cells by lipofection. Prior to delivery, RNP complexes were formed by incubating 37.5 pmol Cas9 with 37.5 pmol sgRNAs per well in Opti-MEM medium at room temperature for 5 minutes. RNP complexes were supplemented with 500 ng EGFP mRNA (Trilink Biotechnologies, L-7201) per well. RNP-EGFP solution was transferred to Lipofectamine CRISPRMAX Cas9 Transfection Reagent (ThermoFisher, CMAX00001) according to manufacturer’s instructions and applied to 6-well plates in drop-wise fashion. After 48 hours, cells were collected and sorted for GFP-positive and DAPI-negative cells using a BD FACSAria III Cell Sorter (BD Biosciences) into 96-well plates at 1 cell per well. Western blotting was used to confirm loss of Cereblon expression, and Sanger sequencing was utilized to verify indel insertion into the target cut site (Forward primer: TGCAGCCTCCACCTTCCAGGTT, Reverse primer: TGTGGTCAGTGTTGAAGCCA).

### Cell-based Tau Aggregation Assay

HEK293T cell lines with inducible GFP-tau were plated at a density of 30,000 cells per well in 96-well black clear-bottom POL-coated plates in DMEM medium supplemented with 10% FBS, 0.1 penicillin/streptomycin, and 10 ng/mL doxycycline to induce GFP-tau expression. After 24 hours, doxycycline was removed by full media change, and treatment with QC-01-175, LDL (Thermofisher, L3486), and tau fibrils were introduced at appropriate concentrations in the new media. After 48 hours, cells were concurrently fixed and permeabilized in 4% paraformaldehyde and 1% Triton X-100 in PBS pH 7.4 at room temperature for 20 minutes. Afterward, cells were washed three times with PBS. After washing, cells were incubated in PBS containing 10 μg/mL Hoescht dye, and plates were sealed prior to imaging. Image acquisition was done using an IN Cell Analyzer 6000 (GE Healthcare Life Sciences). Each well was imaged in a 9 x 9 grid at 20X magnification, covering ≈ 65% of the total well surface area. The resulting images were analyzed using the IN Cell Developer toolbox to quantify the number of Hoescht-positive nuclei and GFP-positive puncta automatically.

### Human Neuronal Cell Culture and Compound Treatment

Approval for working with MGH Frontotemporal Disorders Unit patient-derived cells from the MGH Neurodegeneration Repository was obtained under an MGB/MGH-approved IRB Protocol (#2010P001611/MGH). As previously described (13, 29), fibroblasts from a behavioral-variant FTD patient carrying an autosomal dominant tau-P301L mutation and a healthy individual were reprogrammed into iPSCs, which were subsequently converted into cortical-enriched neural progenitor cells (NPCs) and differentiated into neuronal cells. NPCs were cultured on polyornithine (Sigma Aldrich, P3655) and laminin (Sigma Aldrich, L2020) coated plates (Celltreat; 229106), with DMEM/F12-B27 media (Gibco,11995) supplemented with 20 ng/mL EGF (Sigma Aldrich; E9644), 20 ng/mL FGF (PeproTech; 100-18B), and 5 μg/mL heparin (Sigma Aldrich, H3149). Neurons were differentiated by growth factor withdrawal and half media change twice weekly for four weeks. Treatments were done in six-well plates on 4-weeks differentiated neurons with 2 mL media volume. One mL conditioned media was removed from the culture, and new one mL media pre-mixed with treatments at 2X target concentration was added and incubated at 37°C for the designated period of time. Treatments with QC-01-175, ZXH-4-130 (MedChemExpress, HY-132857), efavirenz (Selleckchem, S4685), and voriconazole (Selleckchem, UK-109496) were dissolved in DMSO before addition to the media.

### Cell Lysate Generation and Western Blotting

Treated neurons in 6-well plates were washed, collected in PBS, and pelleted by centrifugation. Neuronal pellets were lysed in RIPA buffer (Boston Bio-Products, BP-115) supplemented with Roche cOmplete, EDTA-free protease inhibitor cocktail (Millipore Sigma, 11836170001) and phosphatase inhibitor cocktail (Sigma, P0001). Cellular debris was separated by centrifugation at 20,000g for 20 minutes. Supernatant concentration was quantified with the Pierce BCA Protein Assay Kit (ThermoFisher, 23225) according to manufacturer instructions. Gel electrophoresis was performed with the Novex NuPage SDS-Page Gel System by running 10 μg of total protein on pre-cast SDS-PAGE gels (Life Technologies, NPO335). Gels were transferred onto PVDF membranes (Millipore Sigma, IPVH00010) using standard procedures, and membranes were blocked in 5% BSA (Sigma Aldrich, A7906) in Tris-buffer saline with 0.5% Tween-20 (TBST) (Boston BioProducts, IBB-181) for one hour after transfer. Primary antibodies, AT8 (ThermoFisher, MN1020, 1:2000 dilution in TBST), CRBN (Abcam, ab244223, 1:1,000 dilution in TBST), TAU5 (Abcam, ab80579, 1:500 dilution in TBST), and GAPDH (Abcam, ab8245. 1:10000 dilution in TBST) was incubated overnight at 4°C followed by corresponding horseradish peroxidase (HRP)-linked anti-mouse (Cell Signaling Technology, 7076V, 1:2000 dilution in TBST) or anti-rabbit (Cell Signaling Technology, 7074V, 1:2000 dilution in TBST) secondary antibody incubation. Blots were developed with SuperSignal West Pico Plus (ThermoFisher, 34579) and exposed to autoradiographic films (LabScientific; XAR ALF 2025). The resulting films were scanned on an Epson Perfection V800 Photo Scanner and protein bands densitometry was measured using ImageJ and normalized to an internal control.

### Statistical Analysis

Data analysis was done using GraphPad Prism 9 (GraphPad Software) and represented as the means ± SD (*p<0.05, **p<0.01, ***p<0.001). Student’s t-tests were used for simple comparisons. Analysis of variance (ANOVA) followed by Tukey’s multiple comparisons tests post hoc were used for multiple group comparisons.

## Results

### Esterified cholesterol accelerates templated formation of recombinant tau fibrils *in vitro*

Thioflavin T (ThT), a fluorescent small molecule that strongly binds to β-sheet-rich tau aggregates (30), was employed to measure the impact of excess cholesterol and cholesterol ester on *in vitro* tau aggregation kinetics. We utilized non-acetylated low-density lipoprotein (LDL) derived from human plasma as a biological carrier for cholesterol and cholesterol esters (CE). LDL contains roughly 8% cholesterol and 42% cholesterol ester (wt/wt) (31), and in its non-acetylated form, it is readily uptaken by vertebrate cells (32). Tau aggregation can be modeled as a nucleation-dependent growth curve that occurs through three phases: the lag phase (dependent on the initial misfolding event), the growth phase (templated aggregation), and the plateau (substrate dependent) (Figure 1A). We utilized heparin to induce aggregation of monomeric recombinant tau, measured as an increase in total fluorescence. In the presence of only heparin, monomeric recombinant tau aggregation had a calculated time for 50% aggregation (T50) of 33.6 hours. However, in the presence of increasing concentrations of low-density lipoprotein (LDL), the rate of tau aggregation increased with calculated T50 for heparin-induced tau aggregation at 33.8, 25.9, and 20.7 hours for 50, 100, and 200 μg/mL LDL, respectively (Figure 1A).

**Figure 1.**
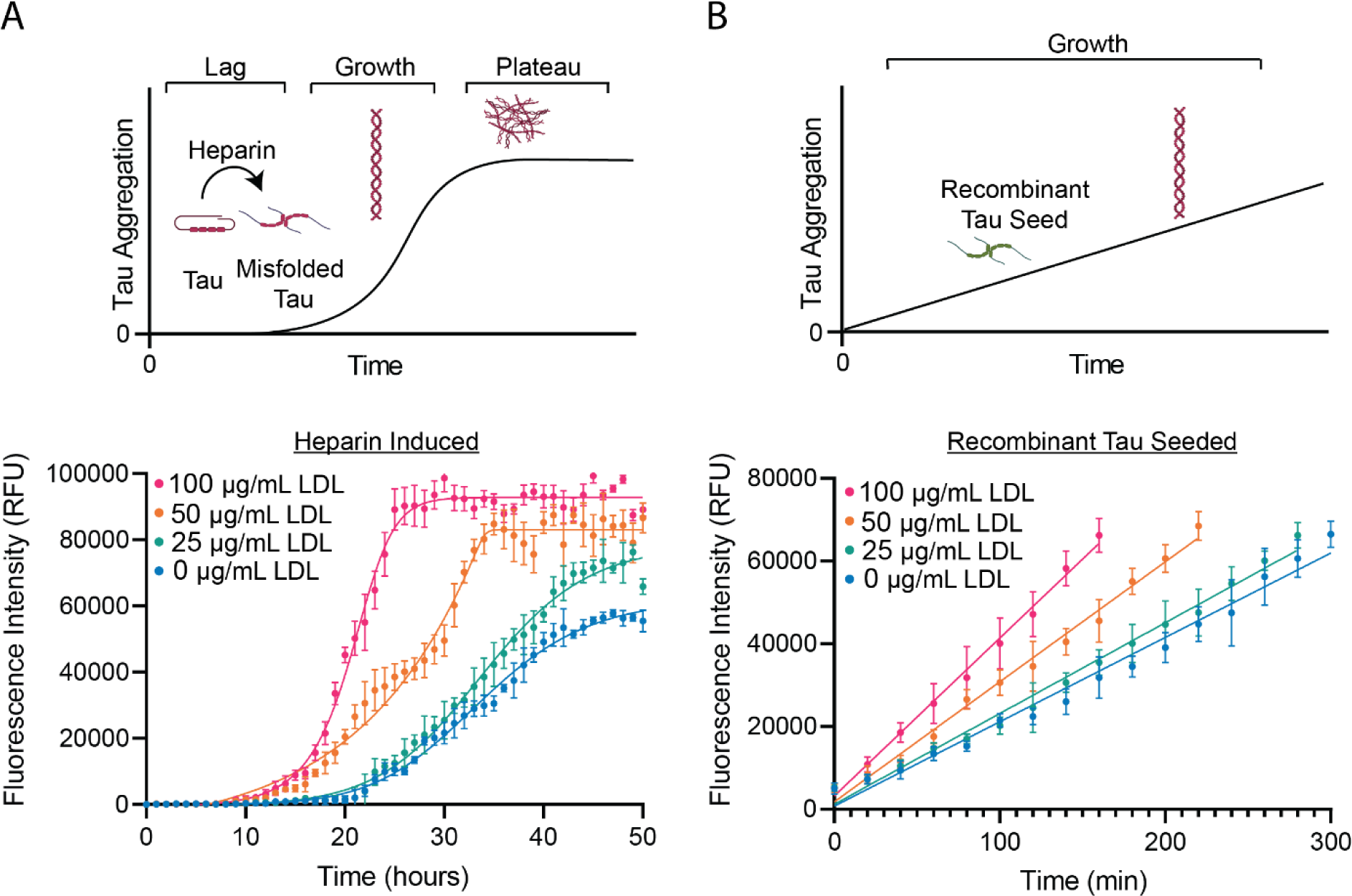
*In vitro* ThT binding assay of recombinant tau in the presence of LDL. A) Aggregation kinetics of recombinant Htau40ΔK280 (20 μM) incubated in Tris pH 7.5 (30 mM), DTT (1 mM), Thioflavin (20 μM), heparin (5 μM) and 0 μg/mL (blue), 25 μg/mL (green), 50 μg/mL (orange), or 100 μg/mL (pink) of LDL. Kinetic curves were fitted as described in *Methods.* Calculated T50 was 33.6 (33.0 to 34.4), 33.8 (33.2 to 34.6), 25.92 (25.3 to 26.5), and 20.7 (20.5 to 20.9) hours for 0, 25, 50, and 100 μg/mL treatment respectively. B. Aggregation kinetics of recombinant tau seeds (1 μg) incubated with recombinant Htau40ΔK280 (20 μM) incubated in Tris pH 7.5 (30 mM), DTT (1 mM), Thioflavin (20 μM), and 0 μg/mL (blue), 25 μg/mL (green), 50 μg/mL (orange), or 100 μg/mL (pink) of LDL. Linear range of assay plotted to measure rate of aggregation as a function of the fluorescence intensity over time. Calculated linear slopes were 204 (194 to 214), 220 (209 to 230), 290 (277 to 304), 382 (356 to 408) for 0, 25, 50, and 100 μg/mL treatment respectively. N = 5 for each individual timepoint, parenthesis represents 95% confidence interval, and error bars are SD.

To account for variations in lag phase duration and potential interactions between LDL and heparin, we conducted the aggregation assay using recombinant tau aggregates instead of heparin to bypass the original misfolding phase. We specifically focused on the ThT assay’s linear range, which was defined as the range between 5,000 and 65,000 relative fluorescence units (RFU). When the reactions were seeded with recombinant tau aggregates, increasing LDL concentration *in vitro* increased the rate of aggregation with calculated linear slopes of 219.5, 290.2, and 381.7 RFU/time for 50, 100, and 200 μg/mL LDL, respectively, versus a slope of 203.9 RFU/time for the control group (Figure 1B). These results show a direct correlation between tau aggregation propensity and LDL cholesterol concentration *in vitro*, suggesting that a direct biological interaction may occur.

### Seed-induced GFP-tau aggregation is exacerbated by LDL in a dose-dependent manner

To measure the impact of cholesterol dysregulation on tau aggregation, we first generated a stable HEK293T line with inducible expression of N-terminal linked GFP-tau. In this tau aggregation sensor cell line, GFP-tau aggregation upon treatment with recombinant tau seeds is detected by the formation of cytoplasmic GFP-positive puncta (Figure 2a, b). We employed a simultaneous fixation and permeabilization step to selectively remove soluble forms of tau, thus increasing the concentration of aggregated species. In the absence of seeds, GFP maintained a diffused cytoplasmic signal. Treatment with increasing concentrations (0.1 μg, 1 μg, and 10 μg) of recombinant tau seeds led to a statistically significant dose-dependent increase of GFP-positive puncta per DAPI-positive nuclei: 1.06±0.32, 6.27±2.35, and 12.8±3.35, respectively relative to vehicle-treated controls (Figure 2b, c). Furthermore, adding recombinant tau seeds to a control HEK293T line expressing only GFP did not cause puncta formation (Supplemental Figure 1).

**Figure 2.**
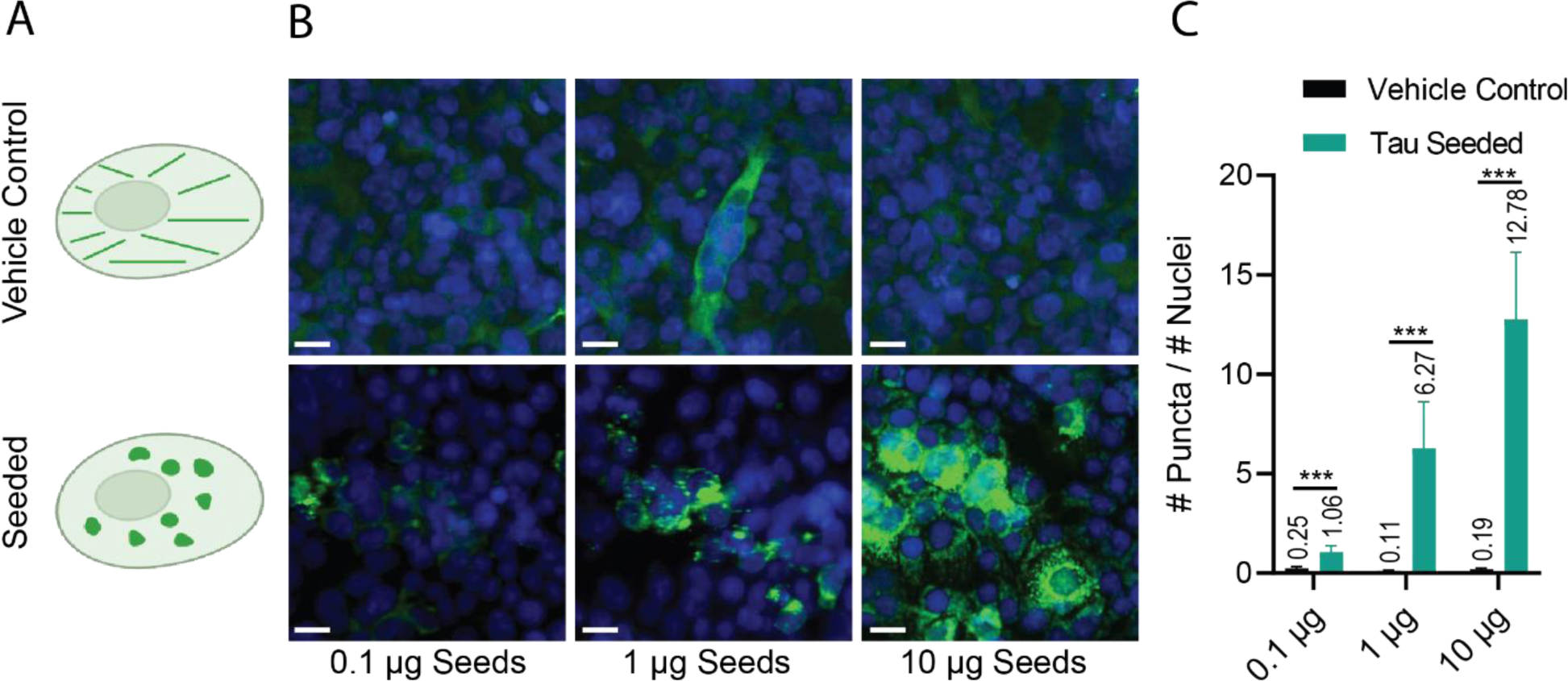
HEK293 GFP-tau Aggregation Assay. A) Schematic representation of non-seeded and seeded GFP-tau. B). Representative images of GFP-tau aggregation assay dose-response to recombinant tau seeds. Nuclei were stained with DAPI (blue) Scale bar is 15 μm. Exogenous tau seeds delivered in lipofectamine 24 hours after GFP-tau induction. Cells were fixed and permeabilized simultaneously 48 hours after seed delivery. C) Quantified dose response of GFP- tau aggregation to recombinant tau seeds. Puncta were automatically quantified using a custom InCell Developer script function and normalized to the number of nuclei detected per image. N = 27 and error bars are SD. *P≤0.05, ***P≤0.001 by unpaired two-tailed Student’s t-test.

To evaluate the impact of cholesterol dysregulation on tau aggregation in a cellular context, we added LDL cholesterol in the presence and absence of recombinant tau seeds to the aggregation sensor line. We added 1 μg of recombinant tau seeds to the aggregation sensor line medium containing either 25 μg/mL, 50 μg/mL, or 100 μg/mL LDL. We observed a trend for the dose-dependent increase in the count of GFP-positive puncta/nuclei of 12.4±1.60, 14.0±2.15, and 19.4±2.54 in LDL + seeds reactions, respectively. Compared to tau seed-only controls, the additional presence of LDL caused a statistically significant (p<0.005) 1.7 to 2.8-fold increase in the count of GFP-positive puncta/nuclei. Furthermore, in the absence of recombinant tau seeds, LDL alone was unable to induce the GFP-tau puncta aggregation phenotype (Figure 3).

**Figure 3.**
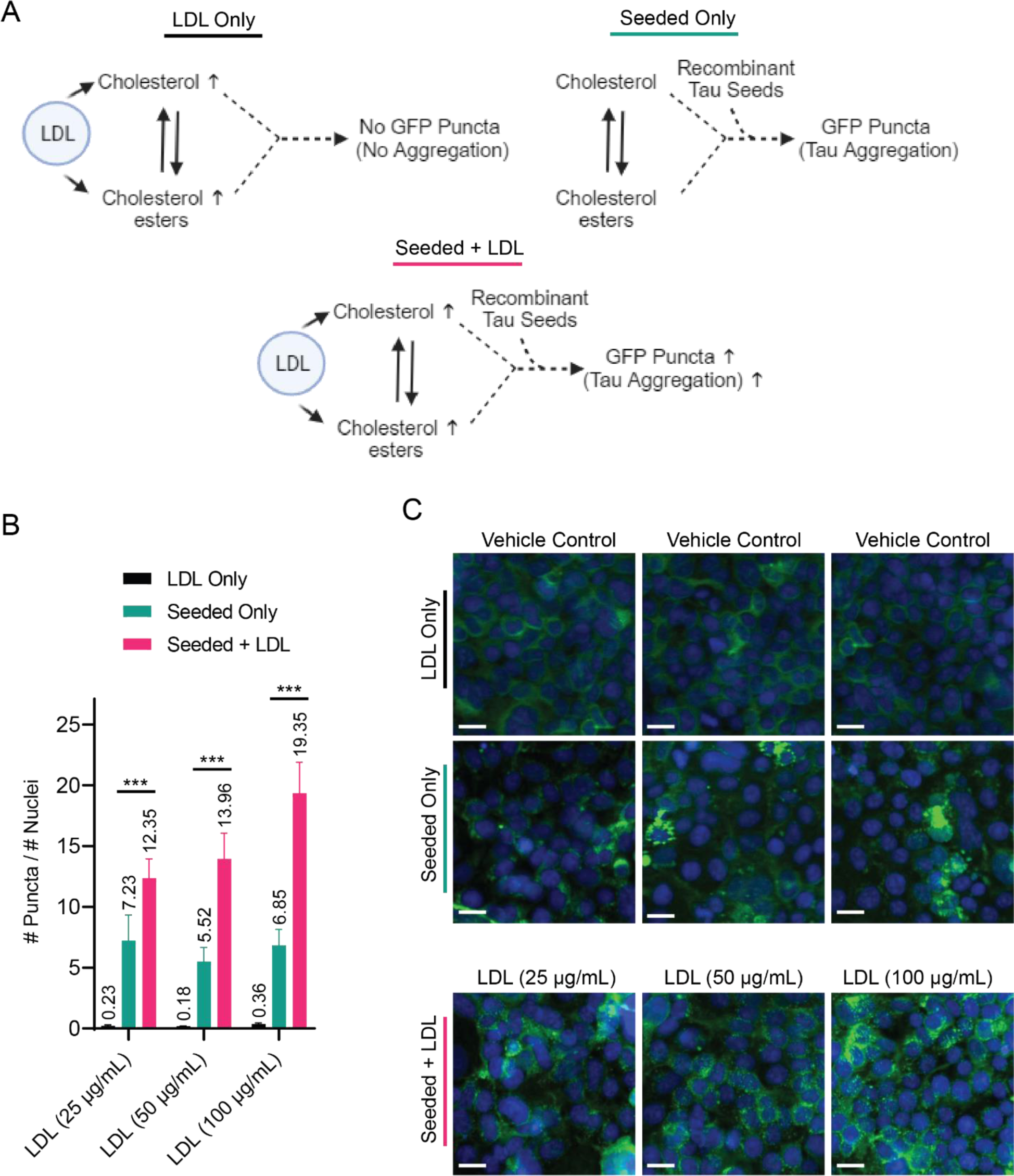
**LDL Enhances Tau Seeding Characteristics in the GFP-tau Seeding Assay**. A) Schematic representation of the pathways targeted by each treatment paradigm. B) Quantified GFP-tau aggregation dose-response to exogenous LDL. LDL and 1ug of exogenous tau seeds were delivered simultaneously 24 hours after GFP-tau induction. Cells were fixed and permeabilized simultaneously 48 hours after seed delivery. Puncta were automatically quantified using a custom InCell Developer script function and normalized to the number of nuclei detected per image. N = 27, error bars are SD. **P≤0.01, ***P ≤0.001 by one-way ANOVA between each of the three groups with post-hoc Bonferroni corrections. C) Representative images from GFP-tau aggregation assay. Nuclei were stained with DAPI

### Degradation of recombinant seeds rescues tau aggregation phenotype

To test if GFP-tau aggregation was dependent on the presence of recombinant tau seeds, we utilized the tau degrader QC-01-175 (33). This heterobifunctional molecule contains a pomalidomide warhead to recruit the CUL4-RBX1-DDB1-CRBN (CRL4^CRBN^) E3 ubiquitin ligase complex (34) and a second warhead to bring the complex proximal to tau species recognized by the clinical tau positron tomography tracer (PET) tracer T807 (flortaucipir) (35). The resulting ternary complex facilitates tau ubiquitination and subsequent proteasomal degradation of the targeted tau species (Figure 4a) (26). Concurrent treatment of the HEK293T GFP-tau sensor cells with QC-01-175 (1 μM) and recombinant tau seeds (1 μM) significantly reduced the GFP-positive puncta/nuclei count from 8.10±1.03 in the seed-only control to 1.80±0.64 in the degrader + seed cells. Furthermore, co-treatment with LDL (50 μg/mL), tau seeds (1 μg), and QC-01-175 (1 μM) rescued the LDL- exacerbated aggregation phenotype (13.1±2.12 puncta/nuclei) to 2.68±1.45 GFP-positive puncta/nuclei (Figure 4a and 4c). As a control for the dependency of QC-01-175 on an interaction with the CRL4^CRBN^ E3 ubiquitin ligase complex, we generated a Cereblon gene (*CRBN*) knockout in the background of the GFP-tau aggregation sensor HEK293T cell line using CRISPR-mediated genome engineering (Supplemental Figure 2). The loss of CRBN rescued the effect of the tau degrader QC-01-175, preventing the degradation of tau species (Figure 4a). Concurrent treatment of *CRBN^-/-^* GFP-tau cells with QC-01-175 (1 μM) and tau seeds (1 μg) caused no significant effect in GFP-positive puncta/nuclei compared to the tau seed-only control. Furthermore, *CRBN^-/-^* GFP-tau cells co-treatment with QC-01-175 (1 μM), LDL (50 μg/mL), and tau seeds (1 μg) also showed no significant difference in the number of tau puncta compared to the LDL (50 μg/mL), and tau seeds (1 μg) only control. Notably, further addition of LDL (50 μg/mL) and tau seeds (1 μg) into *CRBN^-/-^* GFP-tau cell media caused a significant increase in GFP-positive puncta/nuclei to 13.11±2.12, compared to the seed-only control of 8.10±1.03 (Figure 4c).

**Figure 4.**
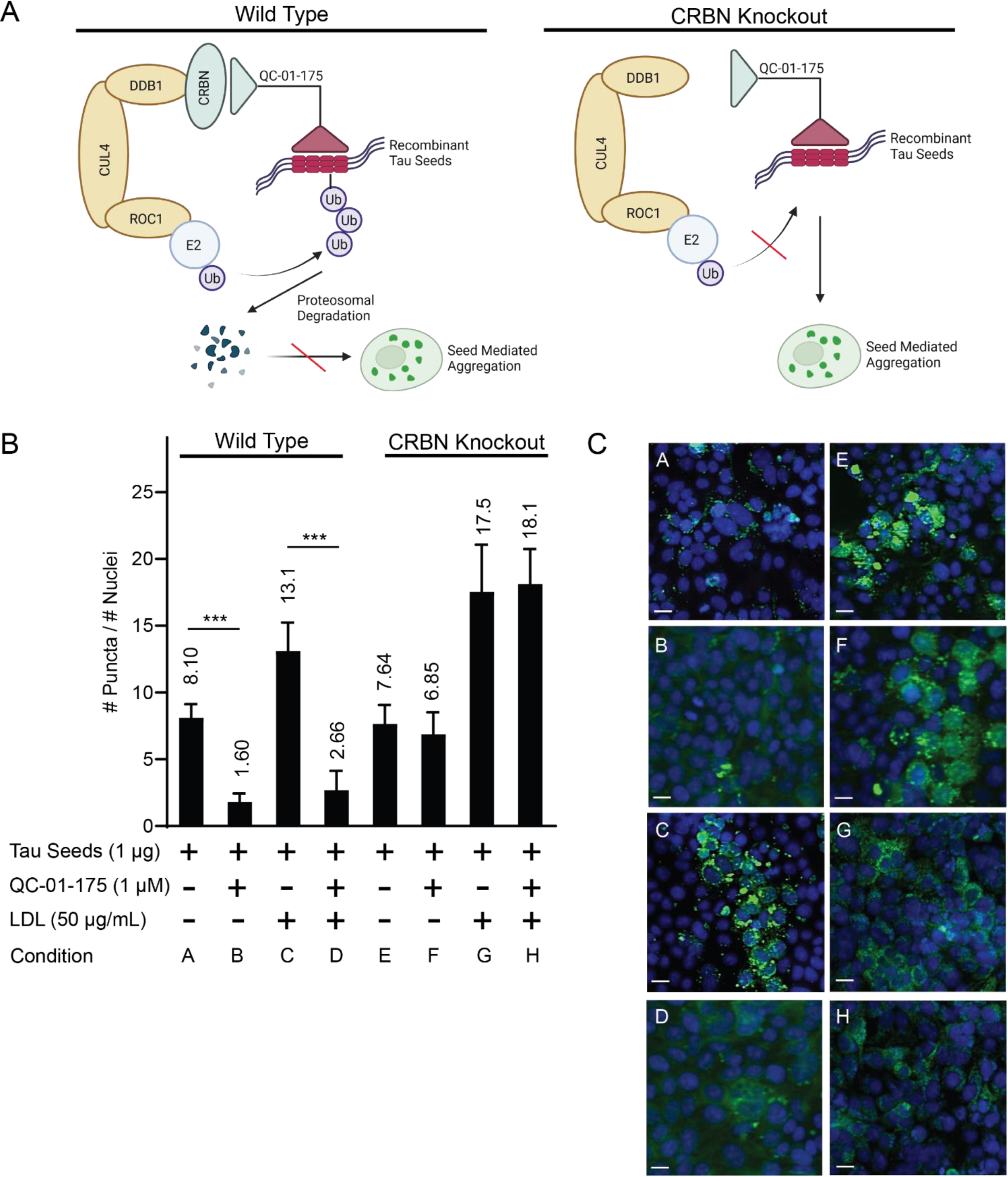
**Degradation of recombinant tau seeds rescues LDL-enhanced aggregation phenotype in a Cereblon-dependent manner**. A) Schematic representation of recombinant tau seed degradation in the absence and presence of CRBN. B) Quantified GFP-tau aggregation assay. CRBN KO and WT GFP-tau sensor lines were treated with a combination of tau degrader QC-01-175 (1 μM), recombinant tau seeds (1 μg), and/or LDL (50 μg/mL) concurrently. ***P ≤0.001 by unpaired two-tailed Student’s t test. N = 27, error bars are SD. C) Representative images from GFP-tau aggregation assay. Images correspond to conditions stated in B. Nuclei were stained with DAPI (blue). Scale bar is 15 μm.

### Cholesterol dysregulation enhances tau accumulation in a human neuronal cell tauopathy model

To investigate the role of tau accumulation and seed-competent tau oligomers in human neurons, we employed patient iPSC-derived neuronal cell models, which are highly relevant for studying neurodegenerative disease phenotypes taking into account the original genetic context and by not requiring overexpression of any transgene (15, 36, 37). To this end, we used iPSC-derived neurons from an FTD patient carrying an autosomal dominant tau-P301L mutation. This patient-specific tauopathy model expresses tau at endogenous levels, and our previous studies have shown that it reproduces disease-relevant tau biochemical and cellular phenotypes (26).

Leveraging the existence of these *ex vivo* tauopathy phenotypes, we sought to validate the use of small-molecule probes of cholesterol metabolism to determine if the increase or reduction of intracellular cholesterol levels affects tau biochemistry in the context of human tauopathy neurons. To this end, we first showed that perturbation of cholesterol turnover via activation or inhibition of Cytochrome P450 46A1 (CYP46A1), the primary metabolizer of neuronal cholesterol (38) (Figure 5a), modulated the ratio of p-tau (pTau S202/T205 detected by the AT8 antibody) to total tau in the iPSC-derived neurons. Treatment of 4-week-old differentiated neurons for 24 hours with efavirenz (0.1 μM or 1 μM), a CYP46A1 activator, led to a reduction (19% and 79% respectively) in the P-tau/total tau levels relative to controls (Figure 5b and 5c). The use of a higher concentration of efavirenz (10 μM) led to a 31% increase in the P-tau/total tau ratio, consistent with previous literature showing the inhibitory effects of efavirenz on CYP46A1 at high concentrations, and thus increased cholesterol (39). Conversely, 24- hour treatment with a potent CYP46A1 inhibitor, voriconazole, resulting in decreased cholesterol turnover, caused a marked 4.20- and 5.89-fold increase in P-tau/total tau relative to controls. Finally, similar to the results with the tau aggregation sensor line, the addition of LDL (50 μg/mL or 100 μg/mL) to the neuronal cultures caused a significant 6.17- and 6.05-fold increase, respectively, in P-tau/total tau (Figure 5B and 5C).

**Figure 5.**
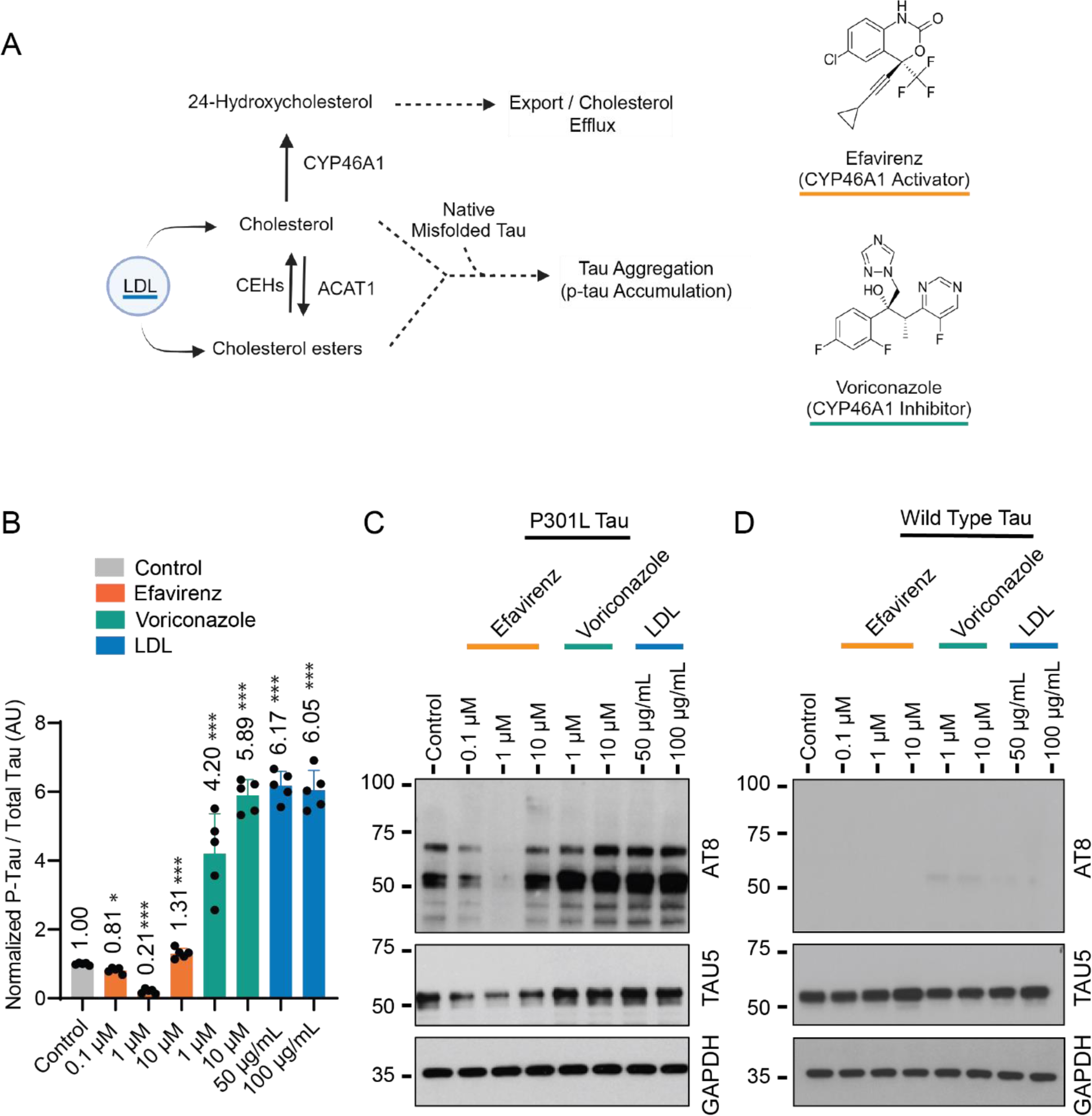
**Cholesterol dysregulation induces pTau accumulation in an iPSC-derived neuronal model harboring a P301L tau mutation**. A) Cholesterol and tau aggregation pathways showing implied mechanism of action by efavirenz, voriconazole, and LDL treatment. B) Semi-quantitative densitometry measurements of the ratio of pTau (AT8) to total tau (TAU5) relative to untreated controls and normalized by loading control GAPDH of 4-week differentiated neurons. Unpaired two-tailed Student’s t test were performed between control and treatment groups. N = 5, error bars are SD. *P≤ 0.05 and ***P≤0.001 by unpaired two-tailed Student’s t test. C) Representative western blot of treatments in 4- week differentiated neurons harboring a P301L tau mutation. D) Representative western blot of treatment in 4-week differentiated neurons derived from a healthy individual (not quantified).

To compare and control for specificity, we utilized an iPSC-derived neuronal cell model from a healthy individual without a genetic predisposition for disease or tau mutations. Changes to cholesterol turnover in the control neurons by activating CYP46A1 with efavirenz (0.1-10 μM) for 24 hours did not result in any measurable increase in the P-tau/total tau compared to vehicle controls. Likewise, inhibiting cholesterol turnover with voriconazole (1 μM or 10 μM) for 24 hours also had no impact on P-tau/total tau. Finally, treatment of control neurons with LDL (50 μg/mL or 100 μg/mL) did not induce a significant change in P-tau/total tau relative to vehicle-treated controls (Figure 5d). Overall, the results in iPSC-derived neuronal models demonstrate that an increase in p-tau caused by cholesterol turnover inhibition or the addition of exogenous LDL occurred exclusively in the tau-P301L neurons.

### Tau degradation can rescue CE-induced tauopathy in a neuronal cell model

To determine if the CE-induced tauopathy is dependent on the presence of misfolded tau in our *ex vivo* neuronal model, we once again utilized the tau degrader QC-01-175, previously shown to reduce p-tau/total tau in the patient iPSC-derived neuronal cell model (26). Treatment of tau-P301L neurons at 4 weeks of differentiation with QC-01-175 (1 µM) for 24 hours led to a 66% reduction in P-tau/total tau ratio relative to the untreated controls, based on the levels of P-tau S202/T205 detected by the AT8 antibody (Figure 6). In agreement with the results in Figure 5, neuronal treatment with LDL (50 µg/mL) led to a 5.89-fold increase in P-tau/total tau relative to vehicle-treated controls. In contrast, co-treatment with LDL (50 µg/mL) and QC-01-175 (1 µM) rescued the P-tau/total tau to basal levels (Figure 6). Conversely, pre-treatment (6 hours) of P301L neurons with ZXH-4-130 (1 µM), a potent heterobifunctional CRBN degrader (Figure 6), followed by co-treatment with QC-01-175 (1 µM) and LDL (50 µg/mL) no longer reduced P-tau/total tau (Figure 6). Altogether, this data establishes a link between misfolded tau phenotypic exacerbation by adding exogenous LDL and provides functional evidence for QC-01-175 effect on tau levels dependency on CRBN levels/activity.

**Figure 6.**
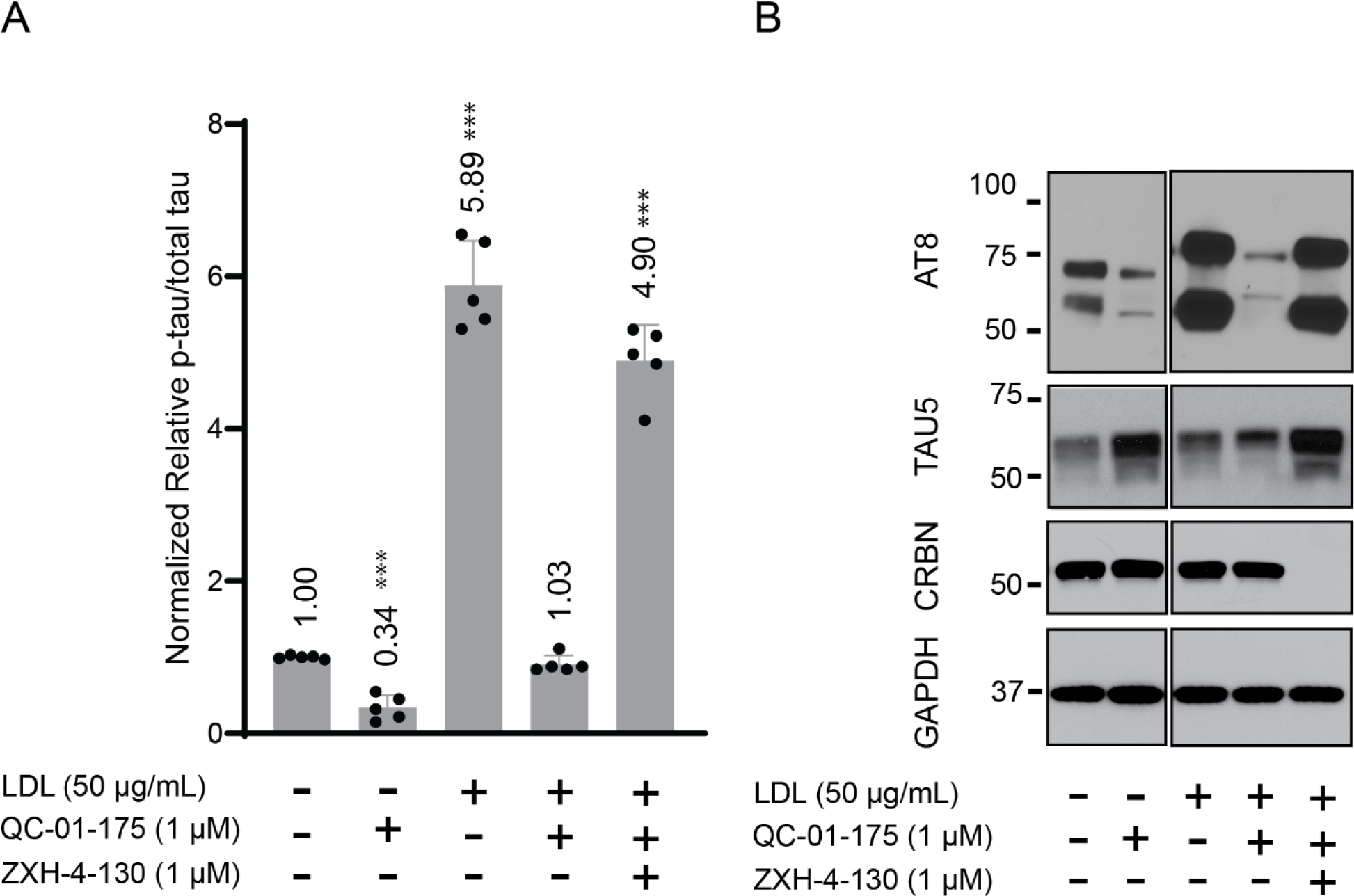
QC-01-175 degrades native misfolded tau and rescues LDL-induced ptau ratios in iPSC-derived P301L tau model. A) Semi-quantitative densitometry measurements of the ratio of p-tau (AT8) to total tau (TAU5) relative to untreated controls and normalized by loading control GAPDH of 4-week differentiated neurons. Treatment was conducted for 24 hours with concurrent treatment with LDL and/or QC-01-175. The treatment group containing ZXH-4-130 were pre-treated with ZXH-4-130 prior to 24- hour treatment with LDL and/or QC-01-175. Unpaired two-tailed Student’s t test were performed between control and treatment groups. *** P≤0.001 by unpaired two-tailed Student’s t test. N = 5, error bars are SD. B) Representative western blot of 24-hour treatment in 4-week differentiated P301L neurons.

## Discussion

Under normal physiological conditions, tau is a highly soluble protein that exhibits little propensity to self-aggregate (40). However, many polyanionic co-factors, such as heparin (41), RNA (42), and fatty acids (43), have been shown to induce tau aggregation *in vitro*. It is postulated that tau aggregation occurs along a nucleation-elongation pathway, whereby stress-inducing detrimental cellular events can lead to tau molecules’ self-association to form an oligomeric seed. Subsequently, other tau molecules join or are recruited by the oligomeric seed, giving rise to higher-order aggregates, protofibrils, and ultimately neurofibrillary tangles observed in tauopathies (44).

Here, we propose a two-hit hypothesis of CE’s ability to accelerate tau aggregation in a seed-dependent manner. *In vitro,* we show that exogenous CE can accelerate the rate of recombinant tau aggregation in the presence of misfolded tau induced by heparin in a dose-dependent manner (Figure 1A). In the absence of a misfolding inducer, LDL can accelerate tau aggregation *in vitro* when in the presence of preformed recombinant tau seeds, also in a dose-dependent manner (Figure 1B). To further test the two-hit hypothesis that CE accelerates tau aggregation in a seed-dependent manner, we employed a GFP-tau aggregation sensor HEK293T line (Figure 2). In this system, LDL alone does not affect tau aggregation, as evidenced by the lack of increased GFP-positive puncta. However, when cells are co-treated with recombinant tau seeds and LDL, we observe a significant increase in GFP-positive puncta compared to cells treated with tau seeds alone (Figure 3). Therefore, LDL enhanced the effect of tau seeds to promote GFP-tau aggregation.

We further measured the impact of exogenous LDL on endogenous tau aggregation in human neurons by utilizing two *ex vivo* iPSC-derived neuronal models, one harboring a tau-P301L mutation and one healthy control. We postulated that tau-P301L neurons would have the appropriate cellular and protein conditions to drive the initial tau misfolding events. Therefore, adding LDL would exacerbate tau misfolding and accumulation, as quantified by the ratio of P-tau to total tau. Of note, despite prolonged differentiation periods, iPSC-derived neurons are relatively immature compared to a human brain, and at endogenous expression levels, tau does not form aggregates nor fibrils. Nonetheless, an early accumulation of P-tau species and high-molecular-weight oligomeric species of reduced solubility correlate with neuronal toxicity in patient-derived neurons but not controls (33). As such, changes to specific P-tau species, such as P-tau S202/T205 (AT8 antibody), are considered as being disease-relevant and were employed in this study as a readout for improvement or worsening of tau accumulation by CE in human *ex vivo* neurons. Indeed, the addition of exogenous LDL caused a significant increase in the P-tau /total tau compared to the untreated control. Conversely, in control neurons, which lacked the cellular environment for tau to misfold, there was no significant difference (or increase or appearance of) in p-tau levels relative to its respective untreated controls (Figure 5d). We orthogonally validate these results utilizing a small-molecule approach with efavirenz and voriconazole, which activate or inhibit cholesterol turnover in the neuronal system, respectively. In addition, this approach allowed us to orthogonally validate the use of LDL as a carrier for cholesterol and CE, as about half its mass can be accounted for by apolipoproteins, phospholipids, and triglycerides (31). In accordance with our earlier results, we showed that inhibition of cholesterol turnover increased P-tau levels only in the P301L tau mutation line, whereas the healthy control line remained unchanged (Figure 5).

It was not surprising that CE alone did not promote the misfolding of monomeric tau, as it lacks the polyanionic charges of common tau fibrillization co-factors. However, fatty acids are exceptionally efficacious drivers of tau fibrillization in physiological conditions, partly due to the anionic head groups and ability to form micelles (43, 45). In fact, evidence from murine models has shown P-tau localization in lipid rafts, suggesting potential interactions between P-tau and membrane lipids (46). Therefore, it is possible that CE, as the conjugated storage form of cholesterol and fatty acids, may directly interact with tau or its misfolded conformers. Furthermore, CEs are stored as lipid droplets, which have been shown to accumulate in neurons in an age-dependent manner and in tauopathies (47). Since misfolded tau can localize along the plasma membranes (48) and reach critical concentrations capable of promoting folding and condensation (49), one hypothesis is that increasing lipid droplet concentration and thereby increasing phospholipid surface areas associated with lipid droplets provided a cellular environment conducive to tau aggregation. Our *in vitro* and *ex vivo* model system studies support a model where the aforementioned environment can significantly increase the rate of tau aggregation.

To further validate the idea that the effect of CE on tau aggregation relies on the presence of seed-competent tau, we employed QC-01-175, a bifunctional tau degrader previously shown to rescue toxicity associated with FTD patient-derived neurons by targeting disease-linked forms of tau, i.e., misfolded species (26). Treatment with QC-01-175 reduced recombinant tau seeds and GFP-tau aggregation in the HEK293 sensor line and promoted the reduction of tau in a tau-P301L patient-derived neuronal model. Notably, this treatment effectively nullified the ability of CE to accelerate tau accumulation/aggregation across the model systems employed. We corroborated these findings through orthogonal methods, with a CRISPR-mediated functional knockout of *CRBN* in the HEK293T sensor line and using a small molecule to induce the degradation of CRBN in the P301L tauopathy model. Both approaches demonstrated that the loss of CRBN abolished the activity of QC-01-175, supporting the idea that proteasomal degradation was the mechanism of tau reduction in both systems (Figures 4 and 6).

It is worth noting that previous research has suggested that CE could directly inhibit proteasome activity in an AD neuronal model (25). Here, the fact that QC-01-175 could degrade P-tau species in an excess CE environment implies that the degrader may function even in a compromised proteasomal environment. This finding has significant therapeutic implications, given that proteasome functionality has been reported to decline with age (50). However, it is also possible that the impact of CE, at different concentrations and on different forms of tauopathy (e.g., AD vs. primary tauopathy) phenotypes may vary, as the consequences of dysregulated cholesterol homeostasis can differ significantly between proteinopathies and brain regions(51).

Altogether, these results suggest that targeting misfolded tau species through proteasome-mediated degradation could be a viable strategy for preventing tau dysfunction caused by cholesterol dysregulation and, more broadly, developing novel therapies for early intervention in age-related neurodegenerative diseases. However, while tau degraders show great therapeutic potential for tauopathies, one concern is that the already hindered proteasome catalytic activity in aged neurons might not be sufficient to degrade enough misfolded- aggregated tau (52). Therefore, it is urgent to investigate multiple avenues of intervention, specifically targeting modifier pathways of tau aggregation, such as cholesterol homeostasis.

## Conclusion

In conclusion, we provide evidence for CE’s ability to accelerate tau aggregation in a seed-dependent manner in an *in vitro* and *ex vivo* system. Furthermore, we showed the rescue of CE-attributed phenotypes through direct degradation of exogenous or endogenous misfolded tau species. In addition, the data provided here suggests that in addition to the ability of heterobifunctional tau degraders to directly lower levels of misfolded tau species, their ability to block intracellular tau seeding and negate the effect of dysregulated cholesterol via proteasomal degradation can potentially be a viable therapeutic option for age-related neurodegenerative disease.

## List of Abbreviations

CE -: Cholesterol esters
CE CRBN: Cereblon
CYP46A1: Cytochrome
P450 46A1 FTD: Frontotemporal dementia
LDL: Low-density lipoprotein
NFTs: Neurofibrillary tangles
PHFs: Paired helical filaments
TBST: Tris-buffer saline with 5% Tween-20
ThT: Thioflavin T

## Declarations

### Ethics approval and consent to participate

Approval for work with non-identifiable human NPC lines was obtained under the Massachusetts General Hospital/MGB-approved Secondary Use IRB Protocol #: 2022P001361, which includes consent for publication of research results.

### Consent for publication

All authors agree to publish.

### Availability of data and materials

The authors declare that the data supporting the findings of this study are available within the paper and its Supplementary Information files. Should any raw data files be needed in another format, they are available from the corresponding author upon reasonable request.

### Competing interests

ML declares no competing interest. S-YK declares no competing interest.

SR serves as a scientific consultant for Sensorium Therapeutics and Entheos Labs.

JEG serves on the SAB of Protego Biopharma, Kaizen Therapeutics, and Contour Therapeutics. He has also received consulting fees from Daiichi Sankyo, Tenaya Therapeutics, Myokardia, Novartis, DiCE, and Rectify Therapeutics.

MCS serves as a scientific consultant for Proximity Therapeutics.

SJH serves on the SAB of Proximity Therapeutics, Psy Therapeutics, Frequency Therapeutics, Souvien Therapeutics, Sensorium Therapeutics, 4M Therapeutics, Ilios Therapeutics, Entheos Labs, and the Kissick Family Foundation FTD Grant Program, none of whom were involved in the present study. SJH. has also received speaking or consulting fees from Amgen, AstraZeneca, Biogen, Merck, Regenacy Pharmaceuticals, Syros Pharmaceuticals, and Juvenescence Life, as well as sponsored research or gift funding from AstraZeneca, JW Pharmaceuticals, Lexicon Pharmaceuticals, Vesigen Therapeutics, Compass Pathways, Atai Life Sciences, and Stealth Biotherapeutics. The funders had no role in the design or content of this article or the decision to submit this review for publication.

### Funding

Funding was provided by the Tau Consortium (Rainwater Charitable Foundation), Alzheimer’s Association- Rainwater Charitable Foundation Tau Pipeline Enabling Program, NINDS/NIH (1R01NS124561-01A1), and the Stuart and Suzanne Steele MGH Research Scholars Program.

### Authors’ contributions

ML, MCS, and SJH participated in the research design. ML, S-YK, and SR conducted experiments. S-YK and JEG contributed research reagents and advice on experimental design and data interpretation. ML performed data analysis. ML, MCS, JEG, and SJH wrote or contributed to the writing and editing of the manuscript with input on the final version by all co-authors.

## Acknowledgments

We thank members of the Haggarty Laboratory and Tau Consortium/Rainwater Charitable Foundation for helpful feedback and discussions. Dr. Eva Mandelkow (German Center for Neurodegenerative Diseases) is graciously thanked for sharing the htau40ΔK280 expression construct used in these studies. Dr. Bradford Dickerson (MGH), Dr. James Gusella (MGH), and Diane Lucente (MGH) are thanked for the generous sharing of patient cell lines.

---

**Supplemental Figure 1.**
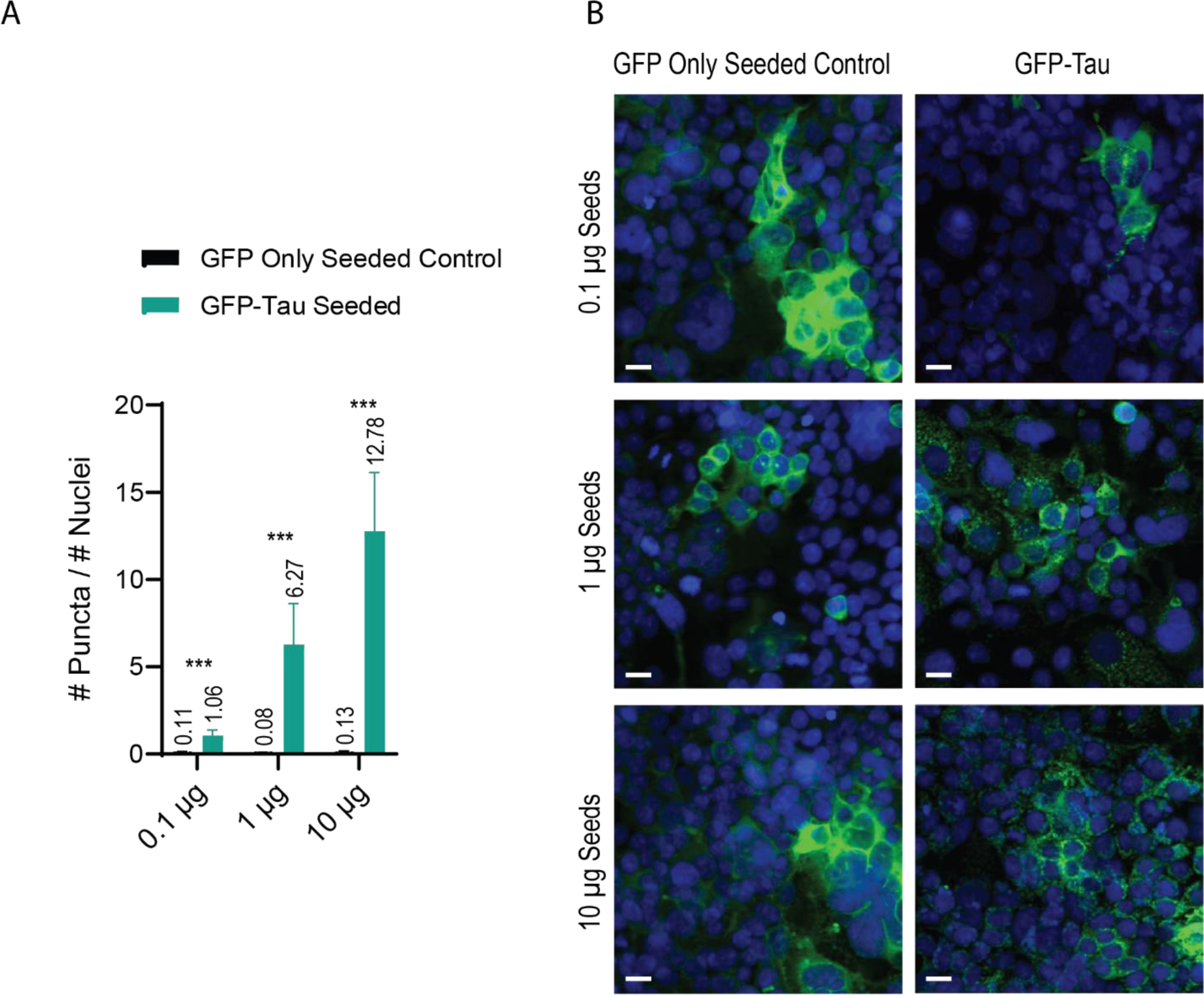
A**d**ditional **HEK293 GFP-tau Aggregation Assay Control.** A) Quantified dose response of HEK293T expressing GFP-tau and GFP only to recombinant tau seeds. Puncta were automatically quantified using a custom InCell Developer script function and normalized to the number of nuclei detected per image. N = 27 and error bars are SD. ***P≤0.001 by unpaired two-tailed Student’s t-test. B) Representative images of GFP-tau and GFP only aggregation assay dose-response to recombinant tau seeds. Nuclei were stained with DAPI (blue) Scale bar is 15 μm. Exogenous tau seeds delivered in lipofectamine 24 hours after GFP-tau induction. Cells were fixed and permeabilized simultaneously 48 hours after seed delivery.

**Supplemental Figure 2.**
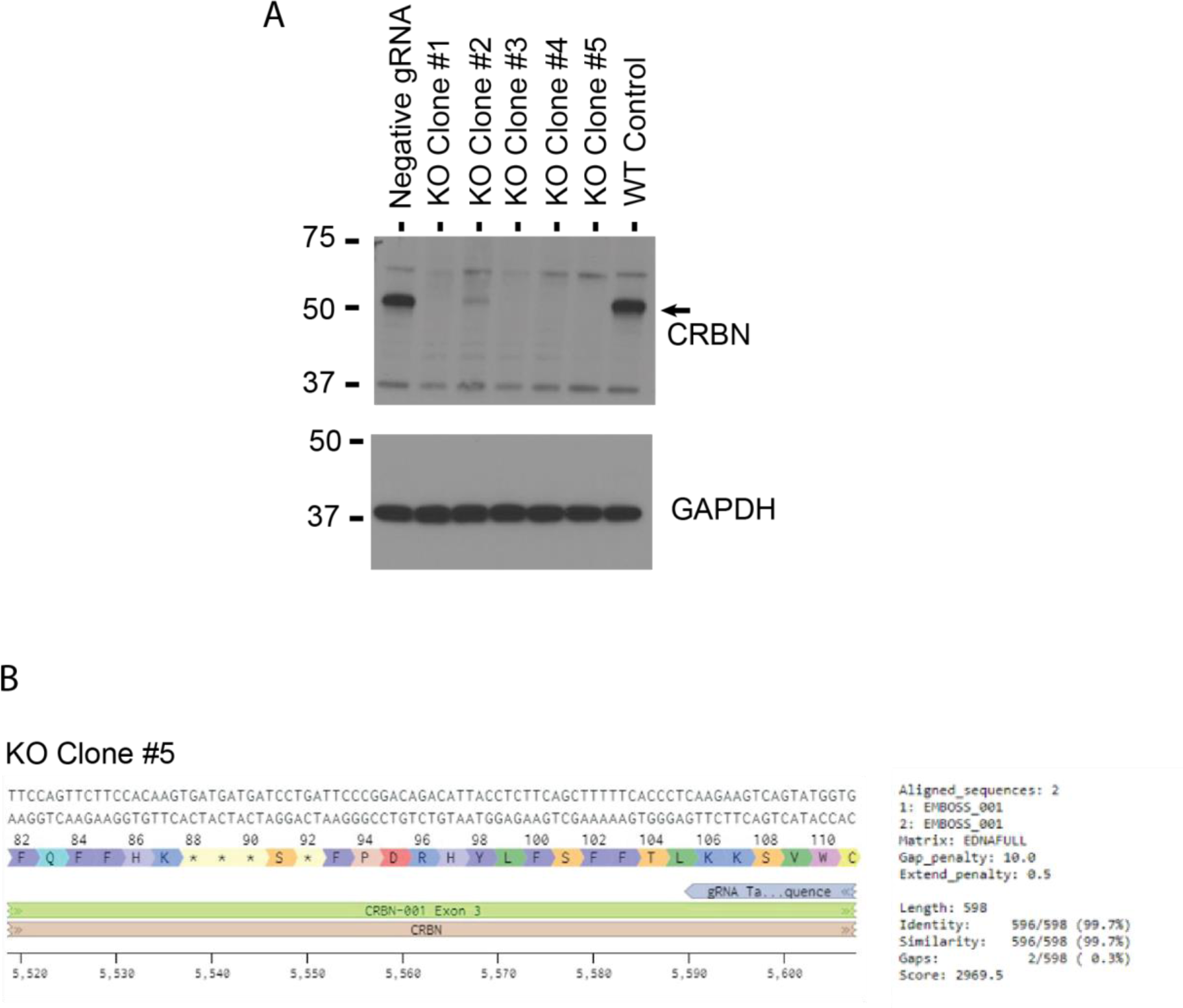
C**e**reblon **Knockout Validation.** A) Confirmational western blot of single-cell sorted clones lacking expression of Cereblon. B) Sanger sequencing alignment for knockout clone #5 around target site.

## References

1. Gigant B, Landrieu I, Fauquant C, Barbier P, Huvent I, Wieruszeski JM, et al. Mechanism of Tau-promoted microtubule assembly as probed by NMR spectroscopy. J Am Chem Soc. 2014;136(36):12615–23.

2. Trinczek B, Ebneth A, Mandelkow EM, Mandelkow E. Tau regulates the attachment/detachment but not the speed of motors in microtubule-dependent transport of single vesicles and organelles. J Cell Sci. 1999;112 (Pt 14):2355–67.

3. Dorostkar MM, Zou C, Blazquez-Llorca L, Herms J. Analyzing dendritic spine pathology in Alzheimer’s disease: problems and opportunities. Acta Neuropathol. 2015;130(1):1–19.

4. Marciniak E, Leboucher A, Caron E, Ahmed T, Tailleux A, Dumont J, et al. Tau deletion promotes brain insulin resistance. J Exp Med. 2017;214(8):2257–69.

5. Lee VM, Goedert M, Trojanowski JQ. Neurodegenerative tauopathies. Annu Rev Neurosci. 2001;24:1121–59.

6. Götz J, Halliday G, Nisbet RM. Molecular Pathogenesis of the Tauopathies. Annu Rev Pathol. 2019;14:239–61.

7. Zempel H, Thies E, Mandelkow E, Mandelkow EM. Abeta oligomers cause localized Ca(2+) elevation, missorting of endogenous Tau into dendrites, Tau phosphorylation, and destruction of microtubules and spines. J Neurosci. 2010;30(36):11938–50.

8. Vaquer-Alicea J, Diamond MI. Propagation of Protein Aggregation in Neurodegenerative Diseases. Annu Rev Biochem. 2019;88:785–810.

9. Clavaguera F, Bolmont T, Crowther RA, Abramowski D, Frank S, Probst A, et al. Transmission and spreading of tauopathy in transgenic mouse brain. Nat Cell Biol. 2009;11(7):909–13.

10. Takeda S, Wegmann S, Cho H, DeVos SL, Commins C, Roe AD, et al. Neuronal uptake and propagation of a rare phosphorylated high-molecular-weight tau derived from Alzheimer’s disease brain. Nat Commun. 2015;6:8490.

11. Ghag G, Bhatt N, Cantu DV, Guerrero-Munoz MJ, Ellsworth A, Sengupta U, et al. Soluble tau aggregates, not large fibrils, are the toxic species that display seeding and cross-seeding behavior. Protein Sci. 2018;27(11):1901–9.

12. Santacruz K, Lewis J, Spires T, Paulson J, Kotilinek L, Ingelsson M, et al. Tau suppression in a neurodegenerative mouse model improves memory function. Science. 2005;309(5733):476-81.

13. Silva MC, Cheng C, Mair W, Almeida S, Fong H, Biswas MHU, et al. Human iPSC-Derived Neuronal Model of Tau- A152T Frontotemporal Dementia Reveals Tau-Mediated Mechanisms of Neuronal Vulnerability. Stem Cell Reports. 2016;7(3):325–40.

14. Silva MC, Nandi GA, Tentarelli S, Gurrell IK, Jamier T, Lucente D, et al. Prolonged tau clearance and stress vulnerability rescue by pharmacological activation of autophagy in tauopathy neurons. Nat Commun. 2020;11(1):3258.

15. Iovino M, Agathou S, González-Rueda A, Del Castillo Velasco-Herrera M, Borroni B, Alberici A, et al. Early maturation and distinct tau pathology in induced pluripotent stem cell-derived neurons from patients with MAPT mutations. Brain. 2015;138(Pt 11):3345–59.

16. Nakamura M, Shiozawa S, Tsuboi D, Amano M, Watanabe H, Maeda S, et al. Pathological Progression Induced by the Frontotemporal Dementia-Associated R406W Tau Mutation in Patient-Derived iPSCs. Stem Cell Reports. 2019;13(4):684–99.

17. Sanders DW, Kaufman SK, DeVos SL, Sharma AM, Mirbaha H, Li A, et al. Distinct tau prion strains propagate in cells and mice and define different tauopathies. Neuron. 2014;82(6):1271–88.

18. Mirbaha H, Chen D, Morazova OA, Ruff KM, Sharma AM, Liu X, et al. Inert and seed-competent tau monomers suggest structural origins of aggregation. Elife. 2018;7.

19. Martinez-Seara H, Róg T, Karttunen M, Vattulainen I, Reigada R. Cholesterol induces specific spatial and orientational order in cholesterol/phospholipid membranes. PLoS One. 2010;5(6):e11162.

20. Aittoniemi J, Róg T, Niemelä P, Pasenkiewicz-Gierula M, Karttunen M, Vattulainen I. Tilt: major factor in sterols’ ordering capability in membranes. J Phys Chem B. 2006;110(51):25562–4.

21. Petrov AM, Kasimov MR, Zefirov AL. Brain Cholesterol Metabolism and Its Defects: Linkage to Neurodegenerative Diseases and Synaptic Dysfunction. Acta Naturae. 2016;8(1):58–73.

22. Chung J, Phukan G, Vergote D, Mohamed A, Maulik M, Stahn M, et al. Endosomal-Lysosomal Cholesterol Sequestration by U18666A Differentially Regulates Amyloid Precursor Protein (APP) Metabolism in Normal and APP-Overexpressing Cells. Mol Cell Biol. 2018;38(11).

23. Linetti A, Fratangeli A, Taverna E, Valnegri P, Francolini M, Cappello V, et al. Cholesterol reduction impairs exocytosis of synaptic vesicles. J Cell Sci. 2010;123(Pt 4):595–605.

24. Liu Q, Trotter J, Zhang J, Peters MM, Cheng H, Bao J, et al. Neuronal LRP1 knockout in adult mice leads to impaired brain lipid metabolism and progressive, age-dependent synapse loss and neurodegeneration. J Neurosci. 2010;30(50):17068–78.

25. van der Kant R, Langness VF, Herrera CM, Williams DA, Fong LK, Leestemaker Y, et al. Cholesterol Metabolism Is a Druggable Axis that Independently Regulates Tau and Amyloid-β in iPSC-Derived Alzheimer’s Disease Neurons. Cell Stem Cell. 2019;24(3):363–75.e9.

26. Silva MC, Ferguson FM, Cai Q, Donovan KA, Nandi G, Patnaik D, et al. Targeted degradation of aberrant tau in frontotemporal dementia patient-derived neuronal cell models. Elife. 2019;8.

27. Sakamoto KM, Kim KB, Kumagai A, Mercurio F, Crews CM, Deshaies RJ. Protacs: chimeric molecules that target proteins to the Skp1-Cullin-F box complex for ubiquitination and degradation. Proc Natl Acad Sci U S A. 2001;98(15):8554–9.

28. Barghorn S, Biernat J, Mandelkow E. Purification of recombinant tau protein and preparation of Alzheimer-paired helical filaments in vitro. Methods Mol Biol. 2005;299:35–51.

29. Zhao WN, Cheng C, Theriault KM, Sheridan SD, Tsai LH, Haggarty SJ. A high-throughput screen for Wnt/β-catenin signaling pathway modulators in human iPSC-derived neural progenitors. J Biomol Screen. 2012;17(9):1252–63.

30. Friedhoff P, Schneider A, Mandelkow EM, Mandelkow E. Rapid assembly of Alzheimer-like paired helical filaments from microtubule-associated protein tau monitored by fluorescence in solution. Biochemistry. 1998;37(28):10223–30.

31. Orlova EV, Sherman MB, Chiu W, Mowri H, Smith LC, Gotto AM, Jr. Three-dimensional structure of low density lipoproteins by electron cryomicroscopy. Proc Natl Acad Sci U S A. 1999;96(15):8420–5.

32. Wang MD, Kiss RS, Franklin V, McBride HM, Whitman SC, Marcel YL. Different cellular traffic of LDL-cholesterol and acetylated LDL-cholesterol leads to distinct reverse cholesterol transport pathways. J Lipid Res. 2007;48(3):633–45.

33. Silva MC, Ferguson FM, Cai Q, Donovan KA, Nandi G, Patnaik D, et al. Targeted degradation of aberrant tau in frontotemporal dementia patient-derived neuronal cell models. eLife. 2019;8:e45457.

34. Zhou L, Xu G. The Ubiquitination-Dependent and -Independent Functions of Cereblon in Cancer and Neurological Diseases. J Mol Biol. 2022;434(5):167457.

35. Dalton RM, Krishnan HS, Parker VS, Catanese MC, Hooker JM. Coevolution of Atomic Resolution and Whole-Brain Imaging for Tau Neurofibrillary Tangles. ACS Chem Neurosci. 2020;11(17):2513–22.

36. Ehrlich M, Hallmann AL, Reinhardt P, Araúzo-Bravo MJ, Korr S, Röpke A, et al. Distinct Neurodegenerative Changes in an Induced Pluripotent Stem Cell Model of Frontotemporal Dementia Linked to Mutant TAU Protein. Stem Cell Reports. 2015;5(1):83–96.

37. Fong H, Wang C, Knoferle J, Walker D, Balestra ME, Tong LM, et al. Genetic correction of tauopathy phenotypes in neurons derived from human induced pluripotent stem cells. Stem Cell Reports. 2013;1(3):226–34.

38. Björkhem I, Lütjohann D, Breuer O, Sakinis A, Wennmalm A. Importance of a novel oxidative mechanism for elimination of brain cholesterol. Turnover of cholesterol and 24(S)-hydroxycholesterol in rat brain as measured with 18O2 techniques in vivo and in vitro. J Biol Chem. 1997;272(48):30178-84.

39. Anderson KW, Mast N, Hudgens JW, Lin JB, Turko IV, Pikuleva IA. Mapping of the Allosteric Site in Cholesterol Hydroxylase CYP46A1 for Efavirenz, a Drug That Stimulates Enzyme Activity. J Biol Chem. 2016;291(22):11876–86.

40. Wille H, Drewes G, Biernat J, Mandelkow EM, Mandelkow E. Alzheimer-like paired helical filaments and antiparallel dimers formed from microtubule-associated protein tau in vitro. J Cell Biol. 1992;118(3):573–84.

41. Goedert M, Jakes R, Spillantini MG, Hasegawa M, Smith MJ, Crowther RA. Assembly of microtubule-associated protein tau into Alzheimer-like filaments induced by sulphated glycosaminoglycans. Nature. 1996;383(6600):550-3.

42. Kampers T, Friedhoff P, Biernat J, Mandelkow EM, Mandelkow E. RNA stimulates aggregation of microtubule-associated protein tau into Alzheimer-like paired helical filaments. FEBS Lett. 1996;399(3):344–9.

43. Wilson DM, Binder LI. Free fatty acids stimulate the polymerization of tau and amyloid beta peptides. In vitro evidence for a common effector of pathogenesis in Alzheimer’s disease. Am J Pathol. 1997;150(6):2181–95.

44. Friedhoff P, von Bergen M, Mandelkow EM, Davies P, Mandelkow E. A nucleated assembly mechanism of Alzheimer paired helical filaments. Proc Natl Acad Sci U S A. 1998;95(26):15712–7.

45. Montgomery KM, Carroll EC, Thwin AC, Quddus AY, Hodges P, Southworth DR, et al. Chemical Features of Polyanions Modulate Tau Aggregation and Conformational States. J Am Chem Soc. 2023;145(7):3926–36.

46. Cutler RG, Kelly J, Storie K, Pedersen WA, Tammara A, Hatanpaa K, et al. Involvement of oxidative stress-induced abnormalities in ceramide and cholesterol metabolism in brain aging and Alzheimer’s disease. Proc Natl Acad Sci U S A. 2004;101(7):2070–5.

47. Glöckner F, Ohm TG. Tau pathology induces intraneuronal cholesterol accumulation. J Neuropathol Exp Neurol. 2014;73(9):846–54.

48. Fanni AM, Vander Zanden CM, Majewska PV, Majewski J, Chi EY. Membrane-mediated fibrillation and toxicity of the tau hexapeptide PHF6. J Biol Chem. 2019;294(42):15304–17.

49. Sallaberry CA, Voss BJ, Majewski J, Biernat J, Mandelkow E, Chi EY, et al. Tau and Membranes: Interactions That Promote Folding and Condensation. Front Cell Dev Biol. 2021;9:725241.

50. Schmidt M, Finley D. Regulation of proteasome activity in health and disease. Biochim Biophys Acta. 2014;1843(1):13–25.

51. Arenas F, Garcia-Ruiz C, Fernandez-Checa JC. Intracellular Cholesterol Trafficking and Impact in Neurodegeneration. Front Mol Neurosci. 2017;10:382.

52. Thibaudeau TA, Anderson RT, Smith DM. A common mechanism of proteasome impairment by neurodegenerative disease-associated oligomers. Nature Communications. 2018;9(1):1097.

